# Pectin cell wall remodeling through PLL12 and callose deposition through polar CALS7 are necessary for long-distance phloem transport

**DOI:** 10.1101/2022.01.29.478312

**Authors:** Lothar Kalmbach, Matthieu Bourdon, Jung-ok Heo, Sofia Otero, Bernhard Blob, Ykä Helariutta

**Affiliations:** Sainsbury Laboratory, University of Cambridge, 47 Bateman Street, Cambridge CB2 1LR United Kingdom; Institute of Biotechnology, University of Helsinki, 00014 Helsinki, Finland

## Abstract

In plants, the phloem distributes photosynthetic products for metabolism and storage over long distances. It relies on specialized cells, the sieve elements, which are enucleated and interconnected through large so-called sieve pores in their adjoining cell walls. Reverse genetics identified *PECTATE LYASE LIKE 12* (*PLL12*) as critical for plant growth and development. Using genetic complementation, we establish that PLL12 enzyme activity is required exclusively late during cell differentiation. Physiological assays suggested a role specifically in phloem long distance transport while short distance symplastic transport does not require *PLL12*. Exploiting CALLOSE SYNTHASE 7 (CALS7) as a polar fluorescent marker, we identified structural defects in sieve pores in *pll12*. Due to the serial arrangement of sieve elements in the phloem, such defects should affect a plant’s ability specifically for symplastic transport over long distances.

## Introduction

In plants, the phloem is a critical system for transport of carbon (*1*), amino acids (*2*), signaling molecules (*3*), and relay of electrical signals (*4*). Bulk transport through the phloem is directional from photosynthetically active source (*i*.*e*., mature leaves) to inactive sink tissues (*i*.*e*., roots, fruits, and buds) (*1*). Sustained disturbance of phloem transport, for example due to pathogen infection, limits plant growth, coordination of developmental processes, and seed yield (*5*). Plants without a phloem cannot develop beyond the early seedling stage (*6*).

The conductive cells in the phloem are the sieve elements (SE), which together constitute the sieve tube. They are adapted to low hydraulic resistance and high pressures (*7, 8*). Key adaptations of SE consist of the selective autolysis of most of their cellular content, including their nucleus (*9*), thickening of the cell wall (*10*), and differentiation of large cytoplasmic connections, the sieve pores in the cell walls between adjacent SEs, the so-called sieve plates (*11*–*13*)

Cell wall remodeling is key to establishing the characteristics of plant tissues during development from a meristematic to a differentiated state. Such modifications play a central role in SE development and function, yet the molecular factors involved are poorly understood. Very recently, a description of the stationary proteome of SE in tobacco identified a suite of cell wall proteins enriched or specific in this tissue (*14*), which however have yet to be functionally characterized. This applies to a lesser degree to CALLOSE SYNTHASE 7 (CALS7), which is required for deposition of the beta-1,3-glucan callose in SE, particularly at the sieve plate (*15, 16*). Yet its subcellular site of activity remains unresolved. Additionally to sieve plate callose, mature SE cell walls contain a dense inner pectin-rich layer (*17*) and display a unique pectin epitope signature, that has been proposed to influence the mechanical properties of the sieve tube (*18*). The underlying tissue-specific mechanisms and their physiological significance, however, have yet to be addressed.

In the primary root, SE first fully differentiate as protophloem SE (PSE) in the meristematic zone close to the root tip while metaphloem SE (MSE) develop later (*19*). The PSE cells are therefore unique among differentiated tissues in their accessibility to high-resolution transcriptomics to describe their entire differentiation process. (*20, 21*). To understand details of SE cell wall modifications during differentiation, we mined such root transcriptomes first for genes enriched or exclusive to sieve elements (*22*). In a second step we cross-referenced such tissue-enriched genes to SE-specific single-cell and bulk RNAseq datasets to identify enrichment late during SE development(*20*). This approach identified *PECTATE LYASE LIKE 12* (*PLL12*) as a critical factor during phloem differentiation. Pectate lyases catalyze the degradation of demethylated pectin into galacturonic acid (*23*). They have been primarily investigated for their role as virulence factors and tissue maceration by necrotrophic pathogens (*24*). However, increasing evidence suggest specific roles for endogenous plant PLLs, for example in accommodating infection threads during symbiosis (*25, 26*) or organ growth and senescence (*27*). Intriguingly, a recent report described the *pll12* mutant to display defects in stomata development and guard cell function (*28*).

Here we provide cell biological, genetic, and physiological evidence for SE-specific pectin degradation through PLL12 required for long-distance phloem transport. Establishing CALS7 as a fluorescent sieve pore marker, we observed sieve plate formation defects in the *pll12* mutant, underscoring a requirement for PLL12 specifically for long-distance phloem transport.

## Results

### *pll12* has delayed root growth defects and is required in differentiating SE

Mining for cell wall modifying enzymes in transcriptomics available at high tissue and temporal resolution for roots and developing SE, (*20, 22*), we identified three *PLL* candidate genes. *PLL26* (*AT1G04680*), *PLL25* (*AT4G13710*) and *PLL12* (*AT5G04310*) were enriched in SE and increasingly abundant as the SE differentiates (**Fig. S1A-C**). Only mutations in the *PLL12* locus, which was the weakest expressed but most enriched for late SE and phloem, showed strong defects in adult plants (**Fig. S1D-E**). Double mutant combinations did not enhance the observed phenotypes (**Fig. S1E**). *pll12* mutants did not show macroscopic defects in early developmental and their seedlings were indistinguishable from the WT until approximately day 8 (**Fig. 1A,B; Fig. S1G**). Then, primary root growth in *pll12* stopped and subsequent lateral root development was initiated but remained greatly reduced compared to WT, leading to a substantially smaller adult root system (**Fig. 1A-B; Fig. S1G-I, Suppl. Video 1**) and a virtual absence of seed production. Complementation with a genomic sequence of *PLL12* under its endogenous promoter rescued all phenotypes (**Fig 1A-B; Fig. S1F-G,I**). Crucially, all mutant phenotypes could equally be restored by expression of the *PLL12* coding sequence under the *NAC45/86-DEPENDENT EXONUCLEASE-DOMAIN PROTEIN 4* (*pNEN4*) promoter, which drove expression exclusively in 1, occasionally 2 SE cells prior to enucleation and cytosolic clearing (**Fig. S2A-D**) (*9*), thus confirming a role for PLL12 specifically in late SE development (**Fig 1A-B; Fig S1G,I**).

**Figure 1:**
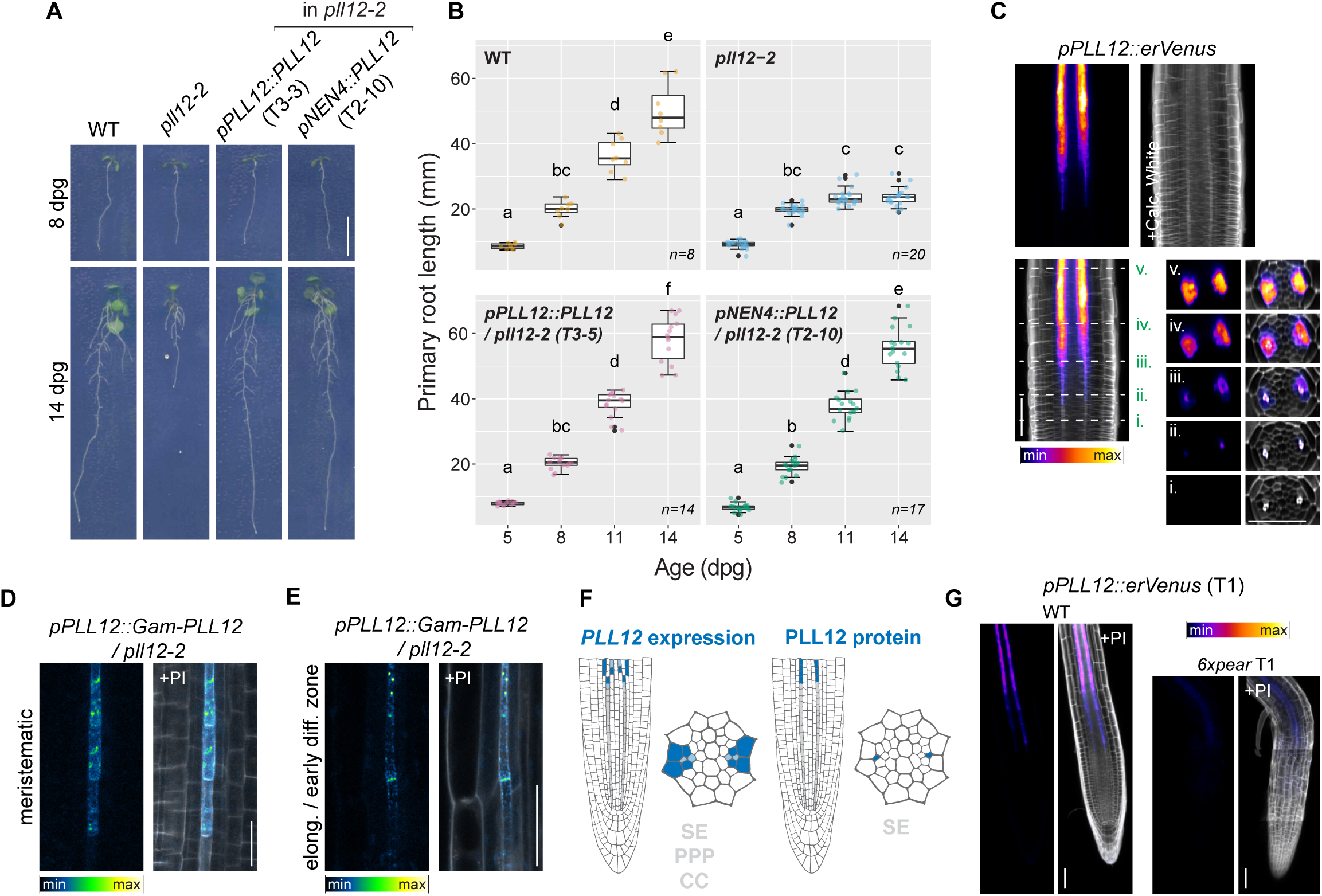
*PLL12* is required in SEs for root growth. **A, B**) Growth and development of *pll12* compared to WT and complemented *pPLL12::PLL12* and *pEN4::PLL12* lines. A) Picture at 8 *vs*. 14 days post germination (dpg) showing *pll12* growth defects absent at 8 dpg but present at 14 dpg; B) Quantification of root length from 5 to 14 dpg (data are presented as box plots with overlaid dot blots, sample size n indicated in each panel; same data is also presented as extended quantification in Figure S1G. **C**) ClearSee preparation of *PLL12::erVenus* reporter line in 5-day-old seedling shows activity in late PSE but strongest in PSE-surrounding tissues; dashed lines and roman numbers indicate positions of cross-sections in right panels. Venus fluorescence shown with as a Look Up Table (LUT) Fire, Calcofluor white (cell walls) is shown in grey. **D, E**) Live-imaging of functional Gam-PLL12 fluorescence in 5-day-old *pll12-2* roots shown as a LUT in Green Fire Blue and cell wall counterstaining with propidium iodide (PI) in grey; Gam-PLL12 is initially diffuse in SE (D) and transiently overlaps with the sieve plate later (E). **F**) Schematic depiction in longitudinal and cross section views of *PLL12* expression domain (left) compared to PLL12 protein (right) in roots. **G**) Representative image of live-imaging of *pPLL12::erVenus* reporter (shown in LUT Fire) in 5-day-old WT (left) and *pear sextuple* mutant (right); cell wall counterstained with PI (in grey), see also Fig. S2. Scale bars represent 10 mm (A), 20 μm (C) and 50 μm (E, G); different letters in (B) indicate significant differences between timepoints and genotypes (*P <* 0.05), Raw data and p-values are presented in **Table S1**.

To validate our initial transcriptome mining data, we sought to obtain a reliable transcriptional reporter by driving a nuclear GFP from the *PLL12* promoter yet were unable to identify T1 individuals with detectable signal. To obtain a robust reporter of *PLL12* expression, we inserted an ER-localized *Venus* coding sequence with terminator between the *PLL12* promoter and the downstream genomic region including its 3’ UTR (*pPLL12::erVen*; **Fig. S2E**), as it had been previously shown for a transcriptional promoter for *ARABIDOPSIS THALIANA MERISTEM L1 LAYER* (*ATML1*) (*29*). This revealed expression in the last PSE before enucleation, which was then followed by increasingly strong expression in the PSE-surrounding tissues of phloem pole pericycle (PPP) and companion cells (CC), which persisted in fully differentiated roots and the aerial vasculature (**Fig. 1C,F; Fig. S2E**). While confirming expression in the SE as initially identified from our transcriptomics, the observed expression throughout the phloem made us wonder how the SE-specific *pNEN4::PLL12* transgene (**Fig. S2A-D**) could fully complement the *pll12* phenotype. In particularly since whole phloem pole single-cell RNA sequencing conducted after our initial transcriptome mining (*21*), perfectly recapitulated our *pPLL12::erVenus* reporter analysis (**Fig S1C**). To determine whether PLL12 protein was SE-specific or across the entire phloem pole, we generated fluorescent protein fusions of *Gamillus-GFP* with the genomic *PLL12* (*Gam-PLL12*) region expressed under its own promoter. Transgenic lines fully complemented the mutant and showed a fluorescent signal specific to the sieve tube. There, Gam-PLL12 initially localized to diffuse peripheral structures in the root meristematic region before it concentrated to discrete foci (**Fig. 1D-F; Fig. S2E**). Later in the SE, PLL12 then briefly localized towards the sieve plate, although never being specific to this structure (**Fig. 1E**).

Phloem differentiation genes should be directly or indirectly under the control of the recently described family of *PHLOEM EARLY DNA-BINDING-WITH-ONE-FINGER* (*PEAR*) transcription factors (*20, 30*). *pPLL12::erVenus* expression is indeed stimulated by inducible overexpression of *PEAR1*, which is among the earliest known organizers of phloem differentiation (*20, 30*) (**Fig. S2G**). To establish a role within the core phloem differentiation program, we transformed *pPLL12::erVenus* into the *pear1 pear2 dof6 tmo6 obp2 hca2* sextuple mutant (*pear sextuple*), which lacks expression of key intermediate and late regulators and transcription factors for phloem and PSE differentiation (*20*). The 23 obtained T1 plants showed either a substantially reduced *pPLL12::erVenus* signal, or no signals at all. In roots with detectable fluorescence, the very weak *pPLL12::erVenus* also appeared no longer confined to the phloem poles (**Fig. 1G**).

### Phloem-specific requirement for pectin degradation through PLL12

Since *PLL12* is expressed in the phloem, required late during SE development, and transcriptionally regulated within the core phloem developmental program, we next wondered about its activity and if the *pll12* phenotype was indeed due to an inability for tissue-specific pectin degradation. While their N- and C-termini differ, the enzymatic domains of pectate lyases are highly conserved from bacteria to land plants and amino acids in the catalytic center are virtually invariable (**Fig. 2A; Fig. S3**) (*31, 32*). We therefore opted for genetic complementation to test for activity of PLL12. In the bacterial pectate lyase PelC of the pathogen *Dickeya dadantii*, which upon infection causes soft rot in plant tissues (*24*), the Arginine at position 218 is part of the catalytic center and required for enzymatic activity (*31, 33*). Substitution of this Arginine to Lysine (R218K) was shown to enzymatically inactivate the protein while not affecting the three-dimensional structure (*34*). We identified the analogous amino acid in PLL12 as R321 (**Fig. S3**). Introducing the R321K mutation into the functional *Gam-PLL12* (*Gam-PLL12*_*R321K*_) transgenes under the control of *pPLL12* and *pNEN4* promoters failed to complement the *pll12* mutant (**Fig. 2B, Fig. S1F**). Importantly, expression and localization of the *pNEN4::Gam-PLL12* transgene, either as WT or inactive version were indistinguishable (**Fig. 2C,D**), suggesting that the *pll12* growth and development defects are indeed due to impaired PLL activity to degrade pectin.

**Figure 2:**
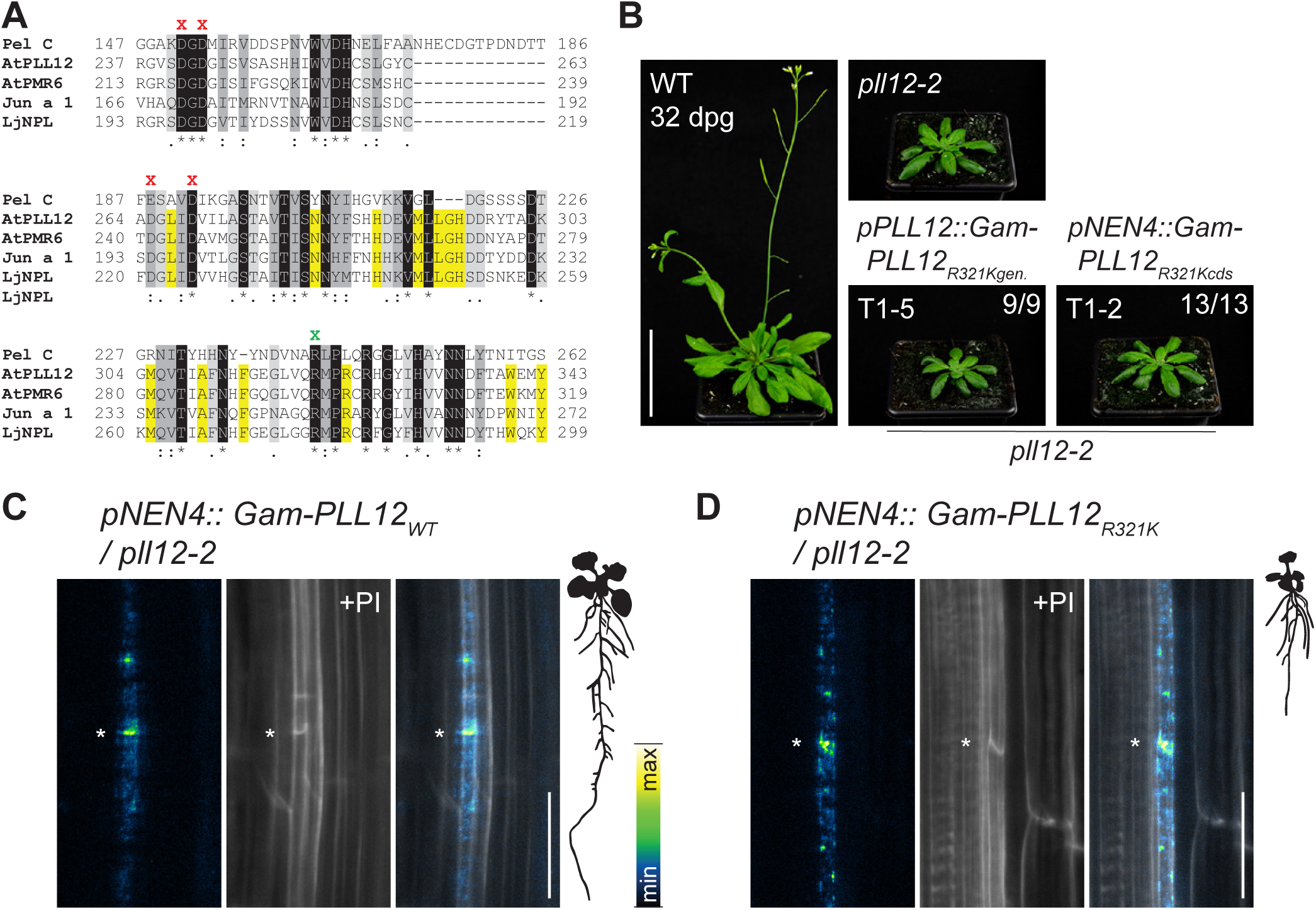
PLL12 pectin-degrading activity is required in SEs. **A**) Partial protein alignment of bacterial pectate lyases PelC and plant PLLs AtPLL12, AtPMR6 (PLL13; closest PLL12 homolog), Jun a 1 (PLL of Juniper with resolved crystal structure), LjNPL (Lotus PLL with inactive point mutation) to identify PLL12 catalytic center; shading indicates similarity; green “X” indicates the invariant Arginine (i.e., R321 in PLL12); for UniProt IDs, references and full alignment see Fig. S3. **B**) Complementation assay in 32 dpg plants with the genomic sequence *pPLL12::Gam-PLL12*_*R312K*_ and the coding sequence *pNEN4::Gam-PLL12*_*R321K*_, both expressed in *pll12-2* mutant background. **C, D**) Protein localization in elongation zone of 5-day-old roots expressing complementing *pNEN4::Gam-PLL12*_*WT*_ (C) and non-complementing *pNEN4::Gam-PLL12*_*R321K*_ (D) in *pll12-2* background, Gam-PLL12 shown in LUT (Green Fire Blue), cell wall counterstained with PI (grey), asterisks indicate sieve plate. Scale bars represent 50 mm (B) and 20 μm (C, D).

Having established that PLL12 enzymatic activity is required in differentiating SE cells, we next wondered if certain types of pectin around the SE might be altered in the *pll12* mutant. To this end, we probed cross sections of the primary root using various pectin antibodies yet failed to observe convincing differences in *pll12* (**Fig. S4A-B**), neither in early differentiating or late differentiated roots. We must bear in mind, however, that other PLLs are still present in differentiating SE of *pll12* (**Fig S1C**) and that therefore net changes on cell wall composition in the *pll12* mutant may well be too subtle to be appreciated in our histological cross sections.

### *PLL12* is required for long-distance but not for short distance cell-to-cell movement

The late appearing but eventually dramatic penalty on growth in *pll12*, associated with pectin degradation defects in the phloem, prompted us to interrogate the physiological basis of its phenotype. In WT, starch synthesized in leaves during the day is degraded to sucrose during the night to be transported into sink tissues. Deficiencies in phloem transport of sucrose are rapidly translated into reduced leaf starch degradation during dark periods (*35, 36*), which can therefore be used as a sensitive proxy for sugar transport defects. We indeed observed significant residual leaf starch at the end of the night in *pll12* but not WT or complemented lines. This was visible as early as 14 days after germination when soil-grown plants (unlike *in vitro* grown seedlings) show hardly any defects in leaf rosette growth and development (**Fig. 3A**). To investigate whether this impaired sugar transport was associated with phloem loading and/or unloading defects, we expressed a cytosolic GFP from the late CC-specific *SUCROSE TRANSPORTER 2* (*pSUC2*) promoter (*37*) in *pll12*. Observation and quantification of fluorescence outside of the phloem in 5-day-old root tips in WT and *pll12* were indistinguishable (**Fig. 3B-C**). This suggests that *pll12* has no fundamental problem with phloem loading, unloading, or plasmodesmata-mediated post-phloem transport. However, since *pSUC2* is expressed in the differentiation zone of the root, it has only limited capacity to instruct us on the efficiency of transport from source tissues to distant sinks. We therefore followed the plant’s ability to transport the symplastic fluorescent tracer 6-Carboxyfluorescein diacetate (CFDA) (*38*) from hypocotyls to the root tip over several days, thus relying on transport over ever increasing distances as the plant grows (**Fig. 3D**). At 5 days (*i*.*e*., prior to root growth defects), *pll12* root tips showed less, though considerable fluorescence compared to WT. However, at 8 days (*i*.*e*., when root growth becomes impaired in *pll12*), no fluorescence was detectable in *pll12* root tips (**Fig. 3D**). When quantified from 4 to 9 days after germination, we observed no difference in phloem transport between genotypes from day 4 to day 6. Yet, over time CFDA transport gradually decreased in *pll12* and virtually no transport occurred from day 8 on. (**Fig. 3E**). In summary, this indicates that *pll12* has no defects in symplastic transport *per se*, as intercellular diffusion over short to medium distances (visualized by free GFP movement in the root tip; **Fig. 3B-C**), is unaffected. Instead, *pll12* has specific defects in long-distance phloem transport, which aggravate as the plant grows (**Fig. 3DE**) and eventually feed back into upstream regulation of source/sink relations in leaves (**Fig. 3A**).

**Figure 3:**
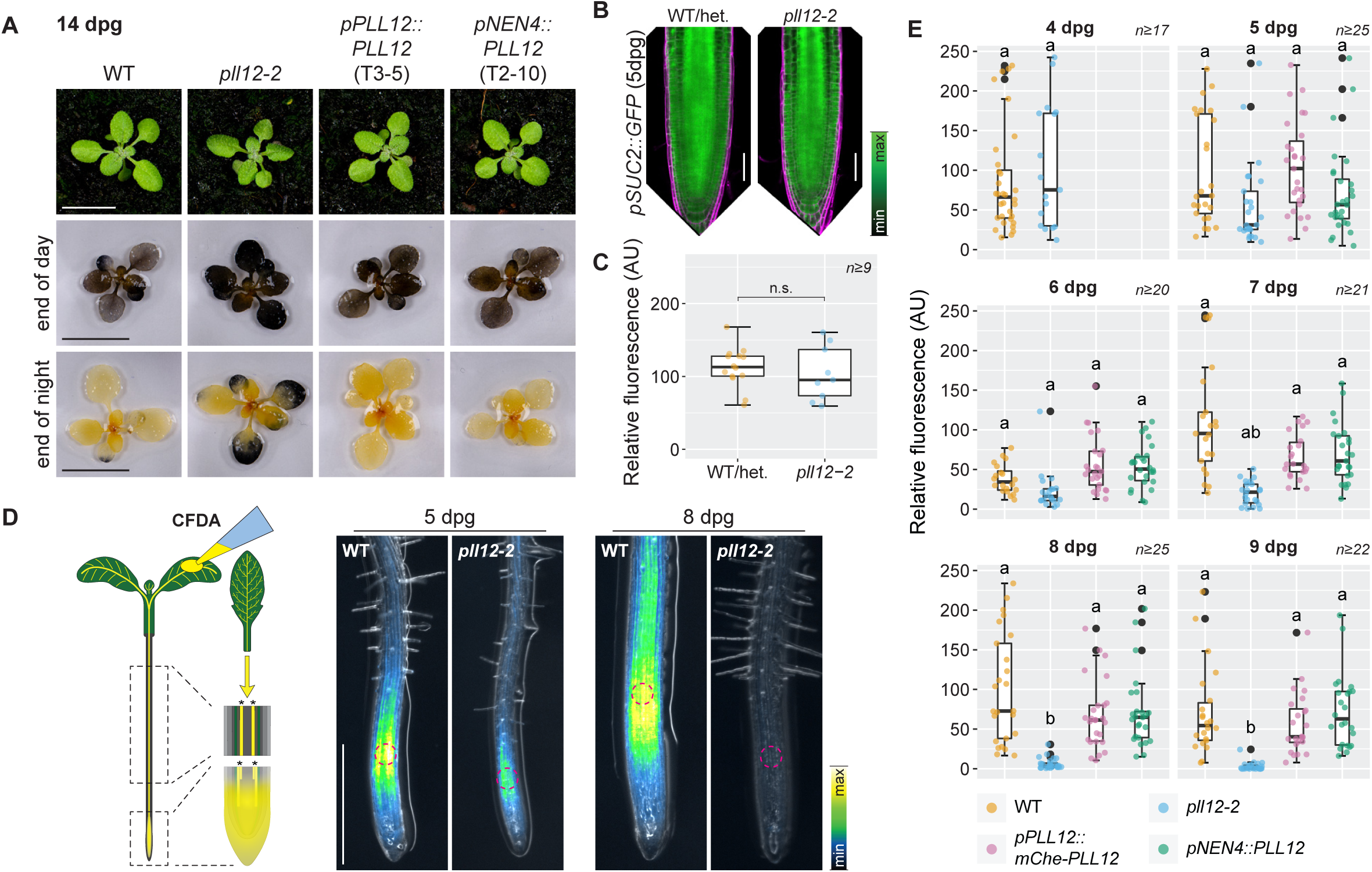
*PLL12* is required specifically for long-distance phloem transport. **A**) 14-day-old plants before staining with Lugol’s iodine solution for leaf starch (upper panels), at after Lugol staining the end of light period (middle panels) and at end of dark period (bottom panels). **B**) Phloem unloading and post-phloem symplastic transport visualized in stable transgenic 5-day-old plants expressing a homozygous single-copy *pSUC2::GFP* (green) in a population segregating for *pll12-2*; cell walls are counterstained with PI (magenta); cytosolic GFP originates from SE, is unloaded and diffuses through the root meristem in both genotypes. **C**) Quantification of unloaded cytosolic GFP of plants from (B) represented as box plots with overlaid dot plot; fluorescence intensity was quantified over a rectangular area 50 μm above the QC; n≥9 roots **D**) Schematic overview of CFDA phloem loading assay and epifluorescence images of root-translocated CFDA approx. 2 h after loading on cotyledons of 5 and 8-day-old plants for WT and *pll12-2*; CFDA fluorescence in LUT Green Fire Blue and overlaid on a dim bright field image for visibility of the root silhouette; note fluorescence decreases in *pll12* but not in WT. **E**) Quantification of fluorescence in root unloading zone approx. 2 h after phloem loading, presented as box plots with overlaid dot plots for WT, *pll12* and *pPLL12::mChe-PLL12* and *pNEN4::PLL12* complemented lines between 4 and 9 dpg; fluorescence intensity scored in region indicated by dashed circle in (D), sample sizes mentioned in each plot. Scale bars represent 10 mm (A), 50 μm (B) and 500 μm (D); different letters in (D) indicate significant differences (*P <* 0.05) between genotypes within one time point. Raw data and p-values are presented for (C) in **Table S3** and for (E) in **Table S4**.

### CALS7 as a fluorescent sieve pore marker

Macroscopic *pll12* growth defects are severe but late and their onset coincides roughly with the emergence of lateral roots that lead to a rapid increase in root system size in WT. This should quickly increase the sink strength of the root system making the growing plant particularly reliant on efficient long-distance phloem transport which *pll12* mutants might be unable to accommodate. A major hydraulic bottleneck for transport through the sieve tube are the sieve pores in the sieve plates between adjacent SEs (*8, 39*). Yet, rapid microscopic investigation of sieve plates is challenging due to their location deep inside tissues, their small size particularly in Arabidopsis (*8*), and lack of bright fluorescent markers. To facilitate and accelerate our microscopic analyses, we sought to establish such as sieve plate specific fluorescent marker by harnessing CALS7. The *cals7* mutant was previously shown to lack deposition of callose specifically in SEs, where callose is most abundant around sieve pores in the sieve plate. Crucially, lack of SE callose leads to reduced phloem transport and stunted growth (*15, 16*), though to a much lesser degree than what we observed in *pll12*. We generated *pCALS7::mNeonGreen-CALS7* plants (*pCALS7::mNG-CALS7*) expressing a fully functional mNG-CALS7 fusion and displaying a strikingly sharp polar localization precisely at the sieve plate in PSE and MSE (**Fig. 4A; Fig. S5A-B**). Closer investigation showed that mNG-CALS7 is initially ubiquitously secreted before it quickly coalesces into a polarly localized domain at the sieve plates 1-2 cells after onset of its expression and on average 3 cells before cytoplasmic clearing (**Fig. 4B,D**). Here, concomitant with its polar localization, CALS7 initiates callose deposition (**Fig. 4C**) approximately 3 cells before cytosolic clearing but clearly after SE cell wall thickening (**Fig. 4D**). This polar localization is susceptible to the endosomal recycling inhibitor Brefeldin A (BFA) (*40*) in early differentiating PSE and MSE but not in fully differentiated SEs later in root development. Combined with the initial, short-lived ubiquitous localization, this suggests that CALS7 polarity relies on endocytosis before the protein is gradually immobilized in a stable plasma membrane domain. (**Fig. S5C**). When expressed ubiquitously with *pUBQ10* promoter, mNG-CALS7 tends to localize polarly in epidermis, cortex, and stele, however neither as sharply defined nor as stable as in SE (**Fig. S5D**). Remarkably, lateral and frontal views directly on differentiated sieve plates revealed that CALS7 is in fact not evenly distributed in sieve plates but lines individual sieve pores between neighboring SEs with a marked donut-like label around the sieve pore necks in 21-day-old major leaf veins (**Fig. 4E**). Altogether, our observations establish CALS7 as a *bona fide* sieve pore protein and corroborate its role for callose deposition during sieve pore formation.

**Figure 4:**
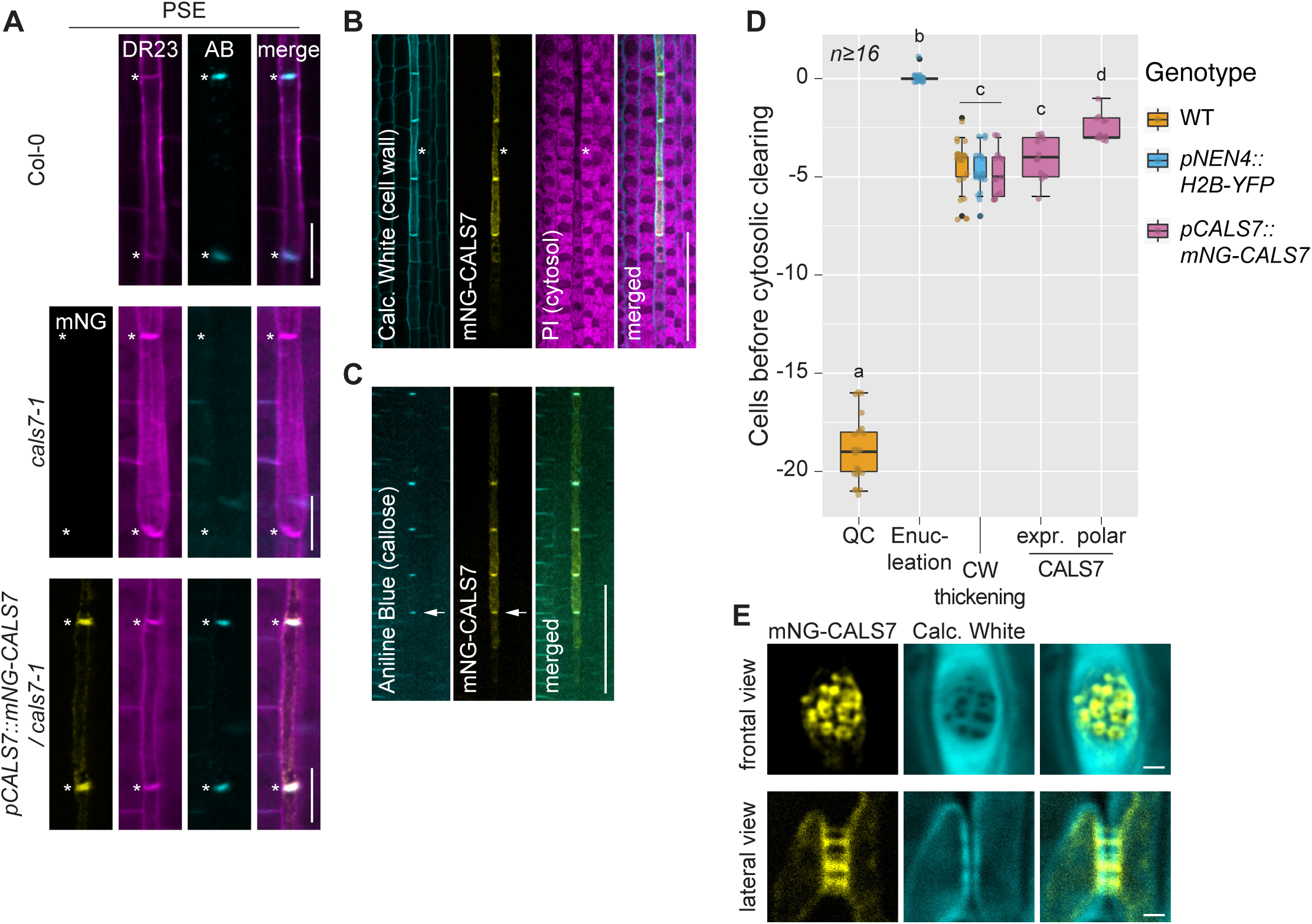
CALS7 is localizing specifically to sieve pores. **A-C**) Co-labeling of ClearSee-treated roots with mNG-CALS7 (yellow) in 5-day-old plants; A) cell wall labeled with Direct Red 23 (DR23, magenta) and Aniline Blue (AB, cyan) to label callose together with mNG-fluorescence in PSE in WT (top), *cals7* (middle) and *cals7* complemented with *pCALS7::mNG-CALS7* (bottom); asterisks indicate position of sieve plates. B) polar mNG-CALS7 localization with respect to cell wall thickening labeled with Calcofluor White (cyan) and PI for cytosolic label upon clearing (magenta); asterisk indicates first cell with degraded cytosol; C) callose deposition (labeled with AB, cyan) coincides with appearance of polar mNG-CALS7. **D**) Quantification of mNG-CALS7 expression and polarization in ClearSee-treated roots, compared to cell wall thickening (visualized with Calcofluor White in all quantified genotypes), enucleation (visualized with nuclear marker *pNEN4::H2B-YFP*), and distance from QC (quantified in Calcofluor White stained WT) in relation to cytosolic clearing; sample size n≥16 PSE cell files. **E**) Frontal and lateral views on differentiated sieve plates in extracted and ClearSee-treated major veins from leaf petioles of 21 dpg plants expressing *mNG-CALS7* in *cals7-1*; mNG-CALS7 labels individual sieve pores (yellow), cell wall counterstained with Calcofluor White (cyan). Scale bars represent 10 μm (A), 50 μm (B, C) and 1 μm (E); different letters (D) indicate significant differences (*P* < 0.05) between measurements. Raw data and p-values are presented in **Table S5**.

### *pll12* has smaller sieve pores

We finally made use of the established CALS7 sieve pore marker to probe whether PLL12 has a role in sieve pore formation or sieve plate patterning to fine-tune SE hydraulic conductivity. Investigating sieve plates in 23-day-old hypocotyls and 21-day-old leaf petioles, we observed that pores in the fully complemented WT-like *pll12* appeared more regular and slightly larger than in non-complemented *pll12* (**Fig. 5A-B**). Additionally, mNG-CALS7 appeared slightly more diffuse at *pll12* sieve plates (**Fig. 5A**). Quantifying sieve pores in secondary tissue of hypocotyls, we noticed considerable variation in pore area in both genotypes. Yet, functionally complemented plants had on average slightly bigger pore areas (**Fig. 5C**). Notably, the largest 25% of sieve pores of *pll12* were smaller than in complemented plants (**Fig. 5D**). The same was observed for pores in the leaf vasculature (**Fig. S6A-B**). These differences appear small. However, it is important to consider that relatively small changes in pore size can have considerable impact on hydraulic conductivity along the entire sieve tube and that individual larger pores contribute most to conductivity across the sieve plate (*8*). The *pll12* transport phenotype may therefore not be due to substantial defects in individual SE but rather a result of moderate effects on individual SE which become severely limiting for long distance transport due to the serial arrangement of SE in the sieve tube.

**Figure 5:**
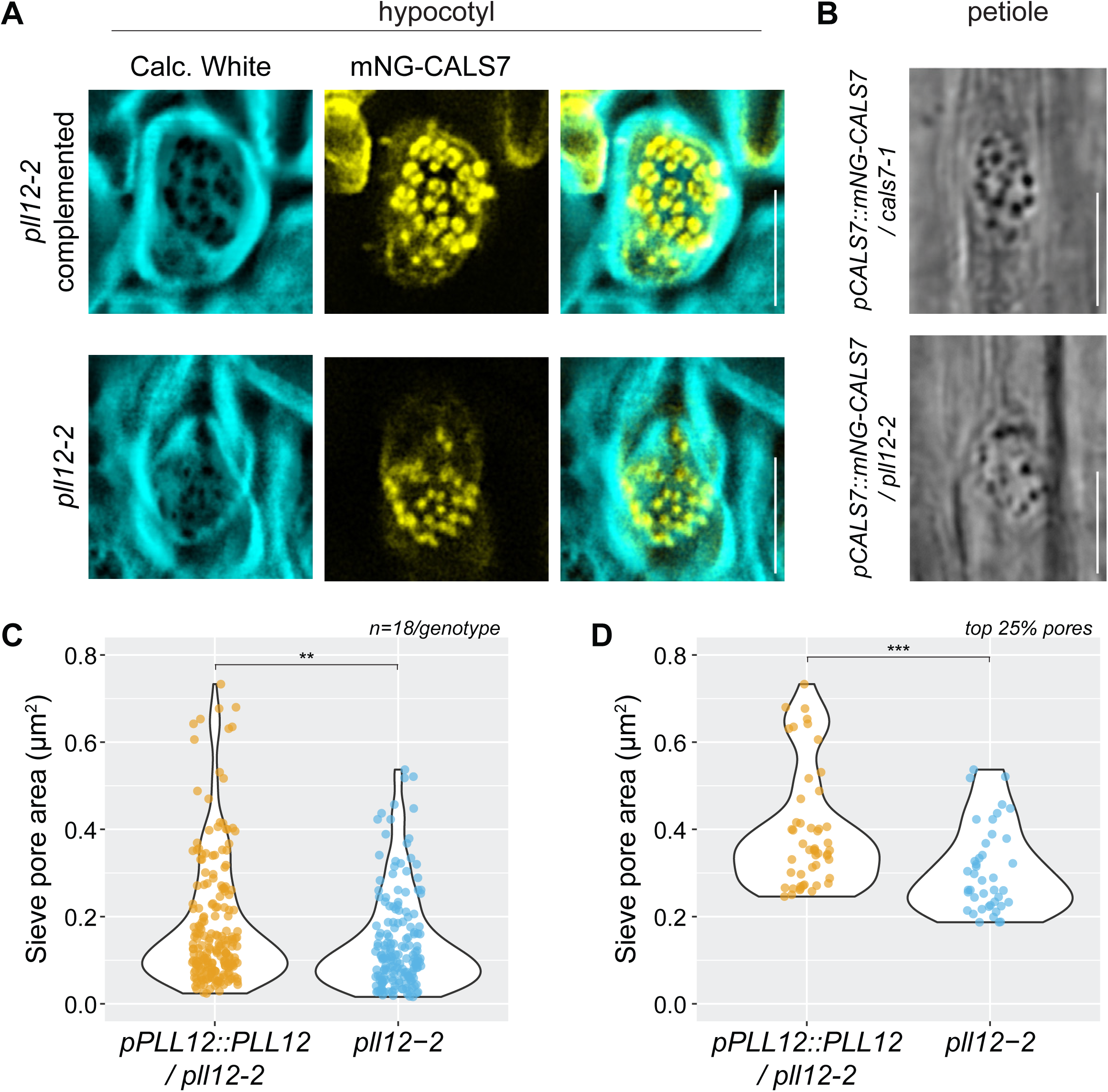
*pll12* has sieve pore defects. (**A**) ClearSee-treated vibratome sections with frontal view on sieve plates of 23 dpg hypocotyls expressing *pCALS7::mNG-CALS7* (yellow) and counterstained with Calcofluor White (cyan). Plants are from a population homozygous for *pll12-2* and *pCALS7::mNG-CALS7* and hemizygous for the complementing *pPLL12::PLL12gen* transgene; complemented plants functionally correspond to WT and are hemi-or homozygous for *pPLL12::PLL12*. **B**) Brightfield image of sieve plates in cleared leaf vasculature of the mNG-CALS7 marker line and *pll12* (carrying a *pCALS7::mNG-CALS7* copy) from 21-day-old leaves. (**C, D**) Quantification of 18 sieve plates per genotype for sieve pore area represented as violin plots with overlaid dot plots considering all (C) and only the largest 25% of each genotype (D); quantified from vibratome sections as presented in (A). Scale bars represent 5 μm (A-B); statistical significance was determined using a Wilcoxon sum rank test, significance levels: ** *= P* < 0.05, *** = *P* < 0.0005. Raw data and p-values are presented in **Table S6**.

## Discussion

Here, we describe *pll12* as a novel phloem mutant, with overall growth and translocation defects that differ fundamentally from previously described mutations affecting this tissue (**Fig. 1A-B, Fig. 3B-E**). *PLL12* appears entirely dispensable for phloem patterning, initiation of SE formation, or generic plasmodesmata-mediated transport. This sets it apart from previously reported mutants such as *octopus* (*ops*) (*41*), *brevis radix* (*brx*) (*42*), *suppressor of max2 1-like 4* (*smxl4) smxl5* double mutants (*43*), or *altered phloem development* (*apl*) (*6*), whose corresponding genes have roles in balancing SE development by antagonizing repressors (*44*), fine-tuning phloem-specific auxin signaling (*45*), or transcriptionally controlling phloem development (*6, 43*). *pll12* mutants are also fundamentally different from previously described general symplastic transport mutants, such as the *gfp-arrested trafficking 1-5* (*gat1-5*) mutants (*46*) or *decreased size exclusion limit 1* (*dse1*) (*47*), which are severely, often lethally affected due to interruptions of any cell-to-cell transport. While such previously described mutants are very instructive in a basic developmental and cell biological context, they offer only limited information on the specific control of long-distance phloem transport throughout a plant’s life.

Although expressed throughout the phloem pole, expressing a *PLL12* transgene cell-specifically shortly before SE enucleation is sufficient for functional complementation thus suggesting a specific requirement of *PLL12* late in SE differentiation (**Fig. 1-3,5**). This is corroborated by the persisting localization of a functional *Gam-PLL12* fusion in the sieve tube when expressed from its endogenous *pPLL12* promoter. A critical function in the sieve tube is also consistent with *pll12* defects in phloem long-distance transport already at the seedling stage. We therefore assume that the pleiotropic defects in adult *pll12* plants are primarily due to global deficiencies in phloem transport, although we cannot formally exclude a role for PLL12 also in other non-vascular tissues, for example as recently reported for guard cells (*28*).

With its defects in transport, adult growth, and seed yield, the *pll12* mutant, although generally more severely affected, resembles *cals7* (*15*). We here confirmed the role of CALS7 for callose deposition around sieve pores and established it as a strong fluorescent sieve pore marker (**Fig. 4**). Like *cals7, pll12* has defects in sieve pore size that are moderate with respect to individual sieve plates. It may appear striking how such minor defects in sieve pore area as observed in *pll12* (**Fig. 5**) can translate into such overwhelming defects in phloem transport and, in consequence, to plant growth. From a theoretical perspective, hydraulic conductivity through a sieve plate is proportional to sieve pore radius to the power of four (*7*). This has two direct consequences. First, relatively small changes in pore size translate into more significant changes in hydraulic flow across a sieve plate. Second, since pore size varies widely within a sieve plate, few relatively large sieve pores contribute by far the most to overall hydraulic conductivity across a sieve plate (*8*). We observed the same heterogeneity in sieve pore size in Arabidopsis, with *pll12* having on average slightly smaller pores. Importantly this difference became more pronounced when we compared only the largest 25% of sieve pores (**Fig. 5B-C**). In consequence, this should lead to *pll12* becoming increasingly unable to keep up with transport demands to its sieve tube.

PLL12 is a pectin degrading enzyme and rendered non-functional upon introduction of the R321K mutation thus providing strong genetic evidence for SE-specific pectin degradation. In its simplest form, lack of pectate lyase activity would increase the amount of de-methylated pectin in the cell wall, which should be detected with the LM19 antibody (*48*). Yet, in the *pll12* mutant, other PLLs are still present in SE, notably *PLL26* and *PLL25* which, based on their expression levels, possibly accumulate at higher levels. De-methylated pectin can equally be processed by polygalacturonases, although involving a different reaction mechanism (*23*). It would therefore not be surprising if changes in pectin composition in *pll12* mutants are very small in quantitative terms, highly localized and were consequently undetectable to us using immunofluorescence.

Since PLL12, although found all around SE cells, briefly localizes to the sieve plate where we indeed observed structural changes, it is tempting to speculate about a role in pectin degradation at the sieve plate. During sieve pore formation cellulose in the primary cell wall is gradually replaced with callose around the necks of the incipient pores, which is then rapidly degraded during sieve pore opening (*11, 12*). This process is reversible and wounding leads to rapid re-deposition of callose (*8, 16*). The enrichment of various other cell wall processing enzymes in fully differentiated sieve tubes (*14*) suggests that cell wall remodeling in mature SE may in fact be constant and dynamic. Pectin is highly abundant in the cell wall (*49*) and de-methylated homogalacturonan, the substrate for PLLs, can effectively mask access to other cell wall polymers for antibody detection (*50*) or enzymatic degradation. Artificial overexpression of pectate lyases in Arabidopsis and poplar increase monosaccharide yields upon saccharification of tissues (*51, 52*). It is therefore conceivable that tissue-specific PLLs can act as gatekeeper to regulate enzymatic modification of non-pectin cell wall polymers by selectively removing homogalacturonan from the cell wall matrix thus granting access to allow localized cell wall modifications to occur and that such processes are interfered with in the *pll12* mutant sieve plates. Future research into endogenous tissue-specific pectin processing is undoubtedly needed and will be crucial to address the complicated inner workings of cell wall remodeling during development, growth, and adaptation.

## Material and Methods

### Plant material and growth conditions

All mutations and transgenic lines are in the Col-0 wild type (WT) background. T-DNA insertion lines used were: *pll26-1* (*SALK_010989*), *pll26-3* (*SALK_147295*), *pll25-1* (*SALK_031335C*), *pll25-2* (*SALK_107510C*), *pll12-1* (*SAIL_1207_A07*), *pll12-2* (*SAIL_1149_C06*), *cals7-1* (*SALK_048921*), *cals7-2* (*SAIL_114_A01*). The mutants *cals7-1* and *cals7-2* (*15, 16*), the pear sextuple mutant as well as the dexamethasone-inducible *pRPS5a::PEAR1-GR* (*30*), and *pNEN4::H2B-YFP* and *pNEN4::erVenus* reporter (*9*) were described elsewhere. Since homozygous *pll12* mutants have severe adult defects and produce almost no seeds, complemented lines hemizygous and segregating for a *pPLL12::PLL12gen*-*pOLE1::OLE1-eGFP* transgene (of pFG7m34GW; see below molecular cloning) were used for bulking, transformation and crossing and then selected against green fluorescence in seeds to obtain pure homozygous *pll12* mutants. For sterile plant culture, seedlings were surface-sterilized either using chlorine gas (by combining 100 ml bleach with 6 ml concentrated HCl for 2-4 h) or in liquid by soaking seeds first in 70% Ethanol + 0.05% Triton-X100 for 3 min followed by soaking in 10% bleach + 0.05% Triton-X100 for 10 min and rinsing seeds subsequently a minimum of 3 times with sterile water. Seedlings were stratified for 2-3 days in dark at 4° C and grown vertically *in vitro* on 50 mL solid half-strength MS growth medium containing 2.2 g/L MS Basal salts (Duchefa M20221) buffered with 250 mg/L 2-morpholinoethanesulfonic acid (MES) and solidified with 0.8% (w/v) Plant Agar (Duchefa P1001). Plants were grown in long-day conditions (16h 22° C / 8 h 18° C) with 150 μmol m^-2^ s^-1^ photosynthetically active radiation (PAR). For prolonged growth *in vitro*, such as for 14 days root growth monitoring, plants were grown on 70 mL solid media and equally spaced at 1.5 cm. For propagation or transformation plants were transferred to compost at 5-10 days post germination. For generation of adult material (phenotyping, Lugol staining, leaf vein extraction) plants were directly sown on compost and stratified for 2-4 days at 5° C in short day conditions (8 h light period with 5 μmol m^-2^ s^-1^ PAR / 16h dark). Plant growth for all adult material was under long day conditions (16h light at 21° C / 8 h dark at 17° C) with 170 μmol m^-2^ s^-1^ PAR during the light period and 65% relative humidity.

### Transcriptome mining

Tissue-specific RNAseq data were filtered for presence (cutoff ≥1.00) of gene expression in S32 transcriptomes (*22*) and in parallel for presence (cutoff ≥1.00) in a bulk transcriptome of protoplasts fluorescence-sorted for the *pCALS7::H2B-YFP* X *pS32::erRFP* marker and representing late differentiating SE (*20*). In a second iteration, the S32 dataset was filtered for enrichment (cutoff ≥2x enriched) compared to whole roots and the *pCALS7::H2B-YFP X pS32::erRFP* transcriptome for enrichment (cutoff ≥2x enriched) compared to meristematic SE cells. Of the 26 PLLs in Arabidopsis, PLL26, PLL25 and PLL12 were present in both enriched datasets. Details about generation and data curation of the mined Transcriptomic are in the respective publications (*20*–*22*).

### Plasmid generation and plant transformation

All Plasmids were generated using the 3x Multisite Gateway® system. For non-destructive selection of transgenes, we established a set of 3x Multisite Gateway compatible destination vectors using the fluorescent seed selection system FAST (*53*). *pFR7m34GW* (FASTred), *pFG7m34GW* (FASTgreen) and pFC7m34GW (FASTcyan) were cloned by generating overlapping PCR products and DNA assembly using NEBuilder. pFR7m34GW was generated by PCR amplifying the pP7m34GW backbone from pH7m34GW from the VIB plant Gateway destination vector collection (*54*) and by PCR amplifying the FASTred cassette (*pOLE1::OLE1gen-TagRFP*) of pFRm43GW. pFG7m34GW was generated by replacing the TagRFP of pFR7m34GW. To this end two overlapping amplicons of the vector backbone were generated and assembled with a third overlapping eGFP sequence. pFC7m34GW was generated by replacing the eGFP of pFG7m34GW. To this end, pFG7m34GW was PCR amplified in two fragments and DNA assembled with a third fragment containing an mTurquoise sequence. The final plasmids were verified by sequencing.

The following entry clones plasmids were already used elsewhere: pENTR_L4-pNEN4-R1 (*9*), pENTR_L4-pCALS7-R1 (*20*). New entry clones were generated by attaching the appropriate attB recombination sites in the PCR primer. To generate entry clones, BP recombination reactions into pDONR-P4-P1r-zeo (1^st^ position), pDONR_P1-P2-kan (2^nd^ position) or pDONR_P2r-P3-zeo (3^rd^ position) were performed according to the manufacturer’s recommendations.

The following entry clones were generated by PCR amplification and direct BP reaction: pENTR_L4-pPLL12-R1 (containing the PLL12 promoter region 1782 bp upstream of the PLL12 start codon), pENTR_L1-PLL12gen-L2 (containing the 3997 bp PLL12 genomic region), pENTR_R2-tPLL12-L2 (containing the 702 bp downstream of the PLL12 genomic region including its endogenous 3’UTR and terminator), pENTR_L1-PLL12cds-L2 (containing the 1554 bp PLL12 coding sequence). pENTR_L4-pSUC2-R1 contains the 1962 bp *SUC2* promoter sequence previously reported (*46*). pENTR_R2-tHSP18.2-L3 contains the 250 bp terminator sequence of AT5G59720 as previously reported (*55*) and was used as terminator in 3^rd^ Gateway position for all constructs except those containing the PLL12 genomic sequence (tagged and untagged) under its own promoter. pENTR_L4-pSUS6-LR1 contains the 2752 bp upstream promoter sequence of *SUCROSE SYNTHASE 6*.

Entry clones for the enzymatically inactive versions of PLL12 (pENTR_L1-Gam-PLL12_R321K_-WT-L2 and pENTR_L1-Gam-PLL12_R321K_-cds-L2) were constructed by first gene-synthesizing short fragments containing the R321K mutation in a genomic (485 bp) or cDNA (461 bp) context; these fragments contained AgeI and EcoRI restriction sites 195 bp upstream and 270 bp downstream of R321 (genomic context) or AgeI and HindIII restriction sites 70 bp upstream and 375 bp downstream of R321 (cDNA context), which allowed for replacing the WT R321 (codon AGG) with the mutant K321 (codon AAG) in pENTR_L1-Gam-PLL12gen-L2 and pENTR_L1-Gam-PLL12cds-L2, respectively, through restriction digestion cloning.

DNA-assembly using the NEBuilder enzyme kit (New England Biolabs E2621) was performed to either modify existing entry clones or clone larger fragments.pENTR_L1-Gam-PLL12gen-L2 fluorescent fusion was generated by DNA assembly of a PCR fragment of the acid-resistant Gamillus GFP (*56, 57*), with matching overlapping 5’ and 3’ regions to be inserted between the endogenous PLL12 N-terminal signal peptide and the start of the PLL12 protein. Gamillus-GFP template for PCR was obtained from a gene-synthesized pENTR-L1-Gamillus-L2. pENTR_L1-Gam-PLL12cds-L2, pENTR_L1-mCherry-PLL12gen; and pENTR_L1-mCherry-PLL12cds were generated accordingly. Constructs for *pll12* complementation tests were recombined into the pFGm34GW destination vector (*pPLL12::PLL12gen-3’UTR*) or the pFR7m34GW destination vector (*pPLL12::Gam-PLL12gen-3’UTR, pPLL12::mCherry-PLL12gen-3’UTR, pNEN4::PLL12cds, pNEN4::Gam-PLL12cds, pPLL12::Gam-PLL12*_*R321K*_*-WT-3’UTR, pNEN4::Gam-PLL12*_*R321K*_*-cds*).

To generate the PLL12 transcriptional reporter in a pseudo-genomic context, a tNos terminator was PCR amplified with overlaps and inserted into a pENTR_L1-erVenus-HDEL-L2 entry clone at the 3’ end of the HDEL ER retention signal by DNA assembly; a pENTR_R2-PLL12gen-3’UTR-L3 clone was generated by PCR amplifying the 4699 bp from the PLL12 ATG to the 3’ end of its putative terminator; eventually, pENTR_L4-pPLL12_R1, pENTR_L1-erVenusHDEL-tNos-L2 and pENTR_R2-PLL12gen-3’UTR-L3 were recombined into pFR7m34GW to the full transcriptional reporter *pPLL12::erVenusHDEL-tNos-PLL12gen-3’UTR*, short *pPLL12::erVenus*.

pENTR_R2-CALS7gen-L3 was generated by DNA assembly of 3 overlapping PCR fragments of the CALS7 genomic region (as a whole including 5’ and 3’ attB recombination sequences) with a PCR-amplified pBluescript II vector backbone; the resulting attB2r-CALS7-attB3 fragment was then recombined into the pDONR_P2r-P3-zeo. To generate pCALS7::mNG-CALS7 and pCALS7::mSc-CALS7, pENTR_L4-pCALS7-R1 and pENTR_R2-CALS7gen-L3 were recombined with either pENTR_L1-mNeonGreen-L2 (into destination vector pK7m34GW) or pENTR_L1-mScarlet-L2 (into destination vector pH7m34GW), which were both donated by Joop Vermeer. Recombination with pENTR_L4-pUBQ10-R1 in pFR7m34GW produced *pUBQ10::mNG-CALS7*. The *pSUC2::GFP* construct was generated by recombining the *pSUC2* entry clone with a pENTR_L1-sGFP2-L2, donated by Marie Barberon, into pB7m34GW. SE-specific plasma membrane marker *pSUS6::Citrine-SYP122* was generated by recombining pENTR_L4-pSUS6-R1 with a pENTR_L1-Citrine-L2 and the plasma membrane protein *SYP122* (At3g52400) (*58*).

Standard molecular practices were used to generate all plasmids, which were verified by sequencing. Expression clones were transformed into electrocompetent *Agrobacterium tumefaciens* cells (strain GV3101) and transformed into Arabidopsis by floral dip (*59*). For each transformation event, at least 15 T1 individuals were screened for expression and/or phenotypes.

### Root growth and biomass assay

Plants were grown *in vitro* and scanned through the agar every day at approximately the same time using an Epson Perfection V700 flatbed scanner at 300dpi. Seedlings were germinated 1.5 cm apart to minimize crowding. Root length was determined by retracing roots in images from the root/hypocotyl junction to the tip using a Wacom Cintiq graphic tablet and measured using Fiji/ImageJ. To measure root dry biomass, plants were germinated and grown on a moist 1:1 mixture of Terragreen pebbles and sand for 42 days. After 14 days, they were supplemented once per week with 1/4 strength Hoagland No.2 media (Sigma Aldrich H2395). For tissue harvesting, substrate was carefully removed from roots under water, the rosette was severed at the hypocotyl and root systems were carefully padded dry before drying for 3 days at 60° C and individually weighted on a precision scale.

### Lugol staining

Staining for leaf starch was performed on 14-day-old plants germinated and grown on compost in long day conditions without elevated light. Plant rosettes for end-of-day samples were harvested in the evening of day 14 approximately 2 h before the end of the photoperiod; rosettes for end-of-night samples were covered with an opaque lid and collected the next morning. Rosettes were immediately transferred into glass jars with 80% (v/v) ethanol and cleared at 55° C for 24h. Tissue was stained with Lugol’s iodine solution (Sigma Aldrich 62650), rinsed twice in pure water, and imaged using a Nikon D5200 DSLR camera.

### CFDA loading assay

Phloem loading was performed essentially as previously described (*60*), with slight modifications. Briefly, a 10x stock was prepared by dissolving 5 mg/ml CFDA (Sigma Aldrich 21879) in Acetone. For loading, 10x CDFA stock was diluted in dH_2_O and 0.5% (v/v) Adigor (Syngenta UK). Plants were grown vertically for 4 to 9 days. For loading, a strip of parafilm was placed above the seedlings on the growth medium and seedlings were carefully pulled upwards to rest with their cotyledons and parts of the hypocotyl on the dry parafilm. Visible transpiration on cotyledons was carefully removed with a fine piece of filter paper, plates were closed and left vertically for 30-60 min so that cotyledons were completely dry. Approx. 0.2 μl CFDA solution was carefully applied on the adaxial surface of a cotyledon using a fine hypodermic needle (26G). Plants were left vertically for 2-3 h to ensure full transport to root tips (which in WT was already the case 20-45 min after loading). Fluorescence intensity was imaged at identical settings for all samples using a Leica M165C stereomicroscope with GFP filter cube and SPOT Advanced imaging software. Images were overlaid with bright field images after acquisition in Fiji/ImageJ and fluorescence intensity was determined as mean intensity values in a 40 pixel circular region halfway between the root tip and emergence of root hairs using Fiji/ImageJ.

### Microscopy and pharmacologic treatments

*pRPS5a::PEAR-GR* was induced by transferring 5-day-old *in vitro* grown plants on half-MS media supplemented with 10 μM dexamethasone for 16 h. 5-day-old seedlings expressing *pCALS7::mSc-CALS7* and plasma membrane marker *pSUS6::mCitrine-SYP122* were transferred into liquid half-MS supplemented with 25 μM BFA or DMSO control for 16 h. Tissue clearing was performed using Clear See and various histological dyes as previously described (*61, 62*). Briefly, tissue was fixed in 4% PFA in PBS supplemented with 0.1% Triton-X100 for 30-60 min, rinsed three times in PBS and transferred to ClearSee solution for clearing in the dark at room temperature for 3-7 days (seedlings) or 7-14 days (hypocotyl vibratome sections and leaf tissue). Direct Red 23 (0.1% w/v), Calcofluor White (0.1% w/v) and Propidium iodide (PI; used in 1:100 dilution of a 1mg/mL stock in dH_2_O) were directly used in ClearSee solution for tissue staining. For aniline blue staining in cleared roots, samples had first to be rehydrated in decreasing concentrations of ClearSee in PBS (90% 80%, 60%, 40%, 20%, 10%, 0%, 0%, 0%) and were then stained with Aniline blue fluorochrome (Biosupplies Cat. No. 100-1) as previously described at 25 μg/mL working concentration in 50 mM Phosphate buffer pH9.5 (*63*). Cell walls of living samples were stained with PI (working solution 10 μg/mL, diluted from a 100x stock in dH_2_O).

Confocal imaging was performed on a Leica SP8 and Zeiss LSM700. Emission and detection settings (in nm) where 405/425-475 (Calcofluor White, Aniline Blue), 488/500-525 (Gamillus, GFP, mNeonGreen), 514/525-550 (YFP, Citrine, and Venus), 552/590-630 (mCherry and mScarlet), 488/600-630 or 552/600-630 (PI and DR23). Images where acquired as stack (transcriptional reporter, quantification of developmental events in Fig. S2B and Fig. 4d, imaging of sieve plates) or as individual slices through the middle of the sieve tube (Gam-PLL12) or the center of the root meristem (*pSUC2::GFP*).

### Confocal image processing and quantification

Images were processed in Fiji/ImageJ, image stacks were z-projected and adjusted for contrast using linear image adjustments over the entire frame or stack. Images of mNG-CALS7 around sieve pores were processed as sub-stacks, and sum-projected. Calcofluor White signal was enhanced for local contrast using the CLAHE plugin. Sieve pores were individually visually identified based on mNG-CALS7 signal and visible pores in Calcofluor White signal. To measure area, pores were retraced using a Wacom Cintiq graphic tablet. To ensure unbiased measurements of areas, genotypes were masked in image file names before measuring and later rematched to genotypes. Cytosolic GFP in root tips was quantified by averaging over a 50 μm x 40 μm rectangular area 50 μm above the QC in Fiji/ImageJ, as described previously (*64*).

### Graphical and statistical analysis

Data analysis used the Tidyverse R package collection and the ggplot2 package for boxplots, dotplots and violin plots. Statistical analysis was performed using the R functions for ANOVA and Tukey HSD tests for multiple comparisons after having determined that parametric tests were applicable using R functions for Bartlett and Shapiro test. Significance values for *P* < 0.05 were grouped using the agricolae package (with alpha = 0.05). Paired significance tests were performed using the Wilcoxon Rank Sum test with significance levels *P* < 0.05 (**) and *P* < 0.005 (***). All specific P-values are in the respective supplementary table.

## Acknowledgment

We thank Hiroyuki Iida and Ari Pekka Mahonen for suggesting the design of the *PLL12* transcriptional reporter, Raymond Wighman and Gareth Evans from the SLCU microscopy facility for technical assistance for imaging and Iris Nieuwland for technical assistance. We thank Caroline Linsdell (Syngenta UK) for supplying a sample of Adigor. Joop Vermeer (*mNeonGreen, mScarlet* entry clones), Niko Geldner (*SYP122* entry clone), Robertas Ursache (pFRm34GW), and Marie Barberon (*sGFP2* entry clone) are thanked for sharing published material.

Alexis Peaucelle, Herman Höfte, Jerome Pelloux, Guillaume Lobet, Paul Dupree, Henry Temple, and all members of the Helariutta Group in Cambridge are thanked for helpful discussions and suggestions. Marie Barberon is thanked for critical reading of the manuscript. This work was funded by the Gatsby Foundation GAT3395/PR3 (Y.H.), ERC Advanced Investigator grant SYMDEV 323052 (Y.H.), Finnish CoE in Molecular Biology of Primary Producers (Academy of Finland CoE program 2014-2019) decision 271832 (Y.H.); University of Helsinki award 799992091 (Y.H.); Swiss National Science Foundation (SNSF) Early Postdoc.Mobility Fellowship project P2LAP3_178062 (L.K.) and a Marie-Curie intra-European fellowship No.: 795250 “SiPoMorph” (L.K.).

## Author contributions

L.K. and Y.H. conceived the project; L.K. performed most of the experiments; M.B., J.O.H., S.O., and B.B. performed additional experiments and provided unpublished material and data; L.K. wrote the manuscript with assistance of Y.H.

---

**Supplementary Figure 1:**
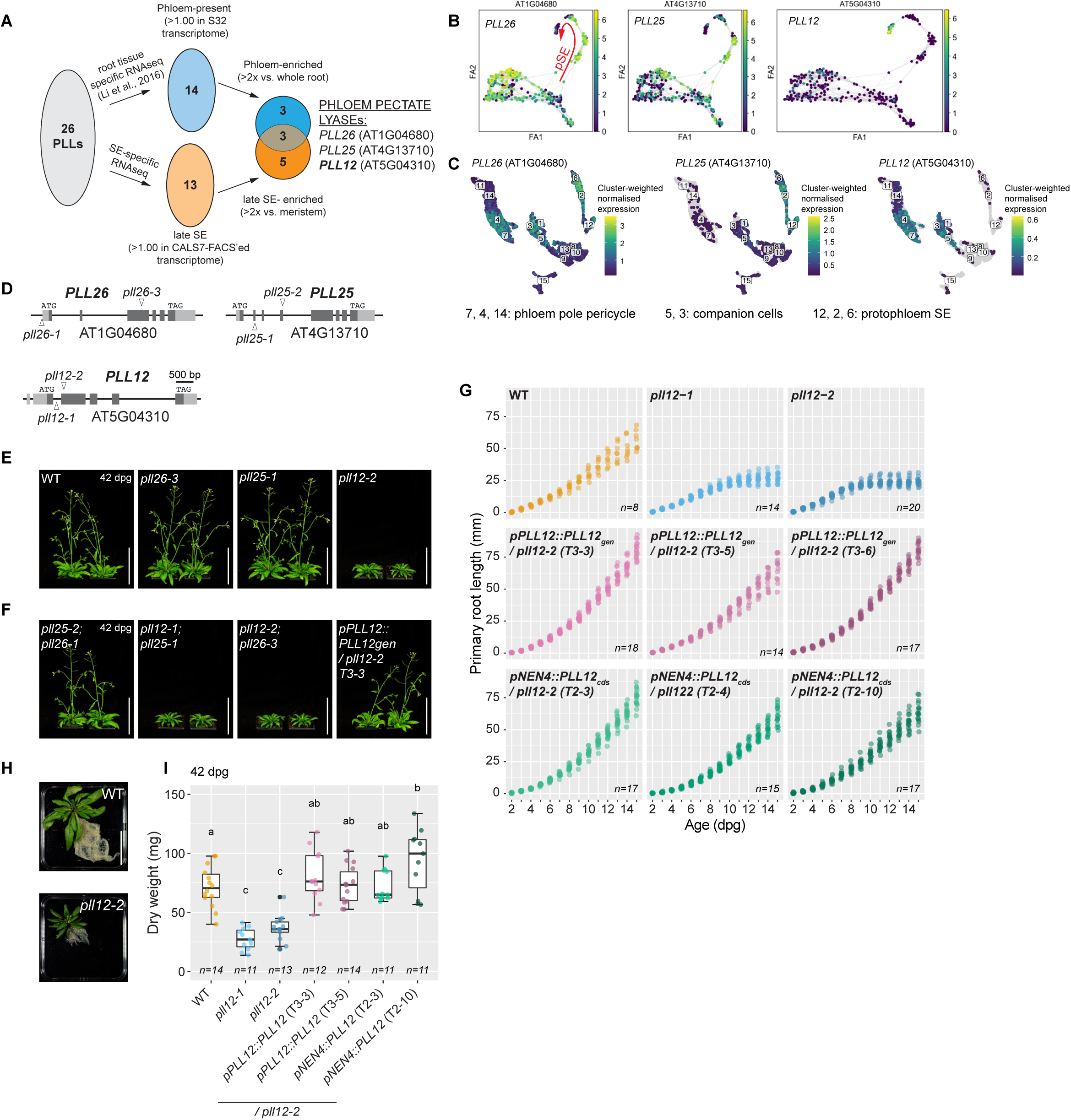
Identification and complementation of the *pll12* mutant. **A**) Of 26 *PLLs* in *Arabidopsis*, 14 are present in SEs of which 6 were enriched (*22*); 13 were expressed in late PSE, of which 8 were enriched and 3 *PLLs* were identified with both approaches. **B**) Forced-direction trajectories show that *PLL12* is most enriched in very late PSE (*20*). **C**) UMAP of whole phloem poles scRNAseq shows abundance of *PLL12* in PSE-surrounding tissues (*21*). **D**) *PLL26, PLL25* and *PLL12* loci and investigated alleles; light grey refers to UTRs, dark grey to exons, triangles indicate location of insertions in mutants. **E**) Phenotypes of *pll* single mutants at 42 days after germination (dpg). **F**) Adult phenotypes of *pll* double mutants and complemented *pll12* at 42 dpg. **G**) Extended root growth phenotyping, including 3 independent complemented lines for endogenous and SE-specific (*pNEN4*) complementation, respectively; source data for WT, *pll12-2, pPLL12::PLL12* (T3-5) and *pNEN4::PLL12* T2-10 at day 5, 8, 11 and 14 is same as in Main Figure 1B. Data corresponds to root length, presented as dot plots from 2 to 15 dpg, sample size n is indicated in each graph. **H**) Root system of sand-grown and extracted adult root systems in WT and *pll12*. **I**) quantification of root dry biomass of 42 dpg sand-grown roots; sample size n indicated in plot. Different letters indicate significant differences *P* < 0.05, raw p-values for significance are presented in **Table S1** for (G) and **Table S2** for (I); scale bars represent 100 mm (E-F) and 50 mm (H).

**Supplementary Figure 2:**
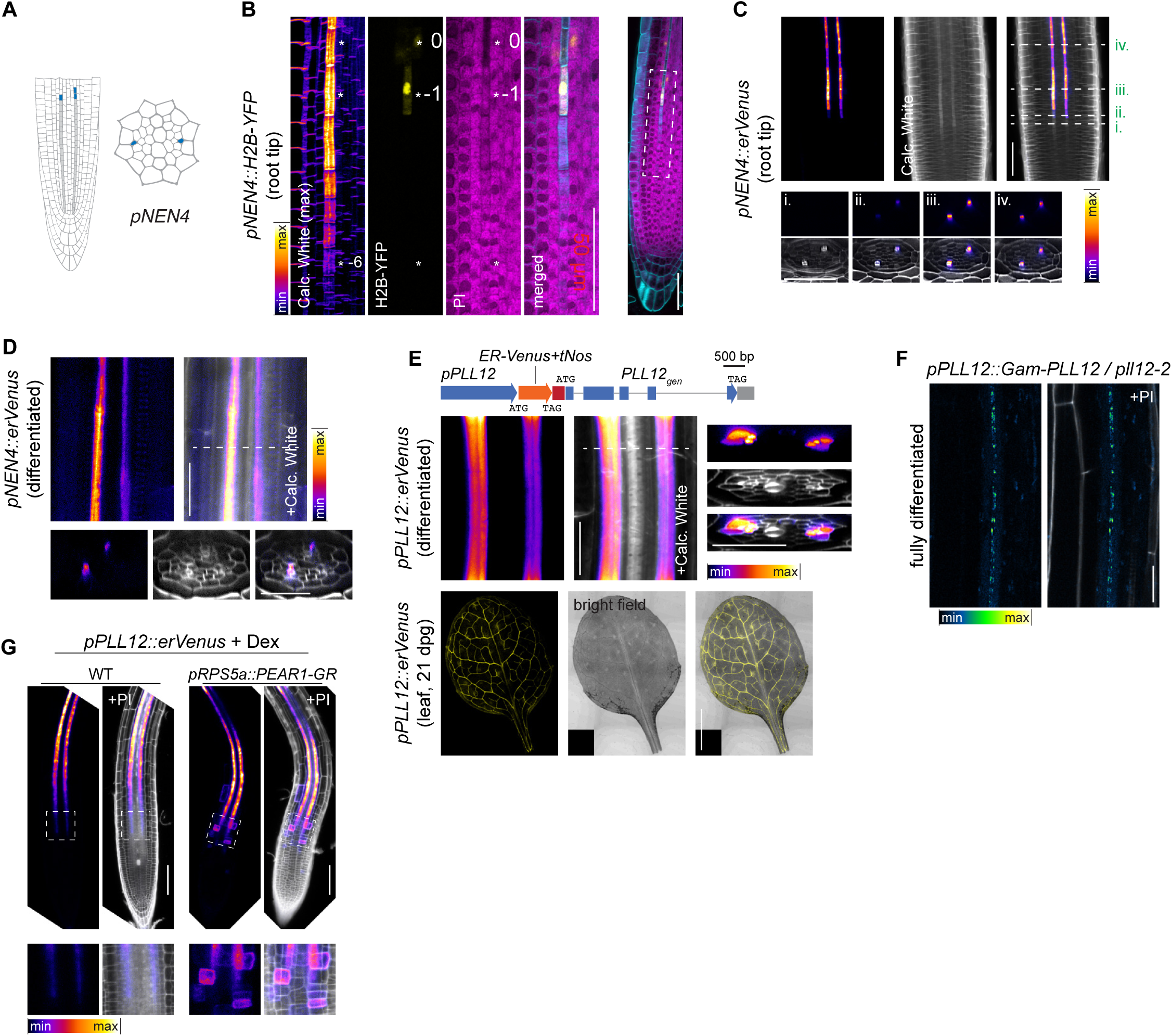
Expression of *NEN4, PLL12* and PLL12 protein localization. **A**) Schematic depiction in longitudinal and cross section views of *pNEN4* promoter activity (in blue; light grey illustrates PSE cell file). **B**) ClearSee and multicolor histological staining of 5-day-old root expressing *pNEN4::H2B-YFP* (in yellow) with Calcofluor White (maximum projection, displayed as LUT fire) and PI (in magenta) to visualize PSE cytosolic clearing; cell “0” corresponds to the first enucleated cell with degraded cytosol; boxed area in right panel indicates position in the root. **C, D**) ClearSee preparation showing *pNEN4::erVenus* transcriptional reporter in early meristematic (C) and late fully differentiated (D) roots in 5-day-old seedling; Venus signal is displayed as LUT fire with Calcofluor White in grey; dashed lines indicate position of cross sections. **E**) *pPLL12::erVenus-3’UTR* transcriptional reporter in a genomic context and expression in fully differentiated and cleared 5-day-old roots (shown with a LUT fire) and cleared leaf of 21-day-old plant (shown in yellow). **F**) Live-imaging of Gam-PLL12 (presented in LUT Green Fire Blue, counterstained with PI in grey) in fully differentiated and functionally complemented 5-day-old *pll12-2* root; cell wall counterstained with PI. **G**) Live-imaging and sum-projection of 6-day-old *pPLL12::erVenus* reporter upon dexamethasone-induction in WT and *PEAR1* dexametasone-inducible overexpressor (*pRPS5a::PEAR1-GR*); close ups in bottom panels are extracted from the boxed area; Venus is presented in LUT fire, cell wall counterstaining with PI in grey. Scale bars represent 50 μm (A, C), 20 μm (D, E, F upper panel), 2 mm (F lower panel) and 100 μm (G).

**Supplementary Figure 3:**
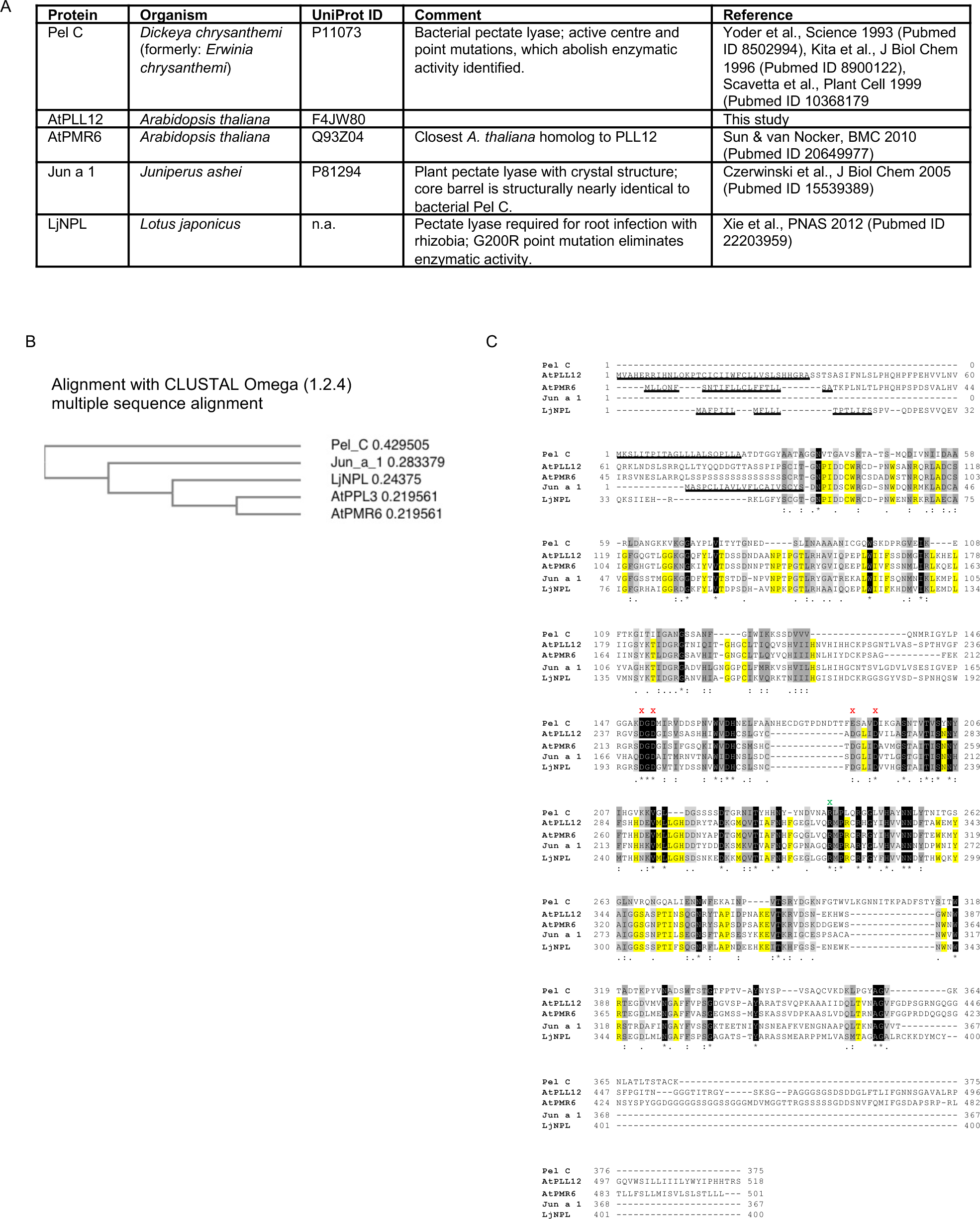
Identification of R321 as the active site of PLL12. **A**) Table summarizing previously described pectate lyases with known amino acid residues essential for activity and/or protein crystal structure. **B**) Protein alignment tree (CLUSTAL Omega 1.2.4). **C**) Alignment of full amino acid sequences using CLUSTAL Omega (1.2.4) multiple sequence alignment. Amino acids shaded grey to black indicate similarity or identity across all pectate lyases; amino acids highlighted in yellow indicate identity between the plant species. Red “X” indicate Ca^2+^ binding sites required for enzymatic activity; green “X” indicates active center of enzyme with reference to Pel C (*33, 34*). N-termina signal peptides are underlined. Note that if signal peptides are included, R218 of Pel C is in fact at position 240.

**Supplementary Figure 4:**
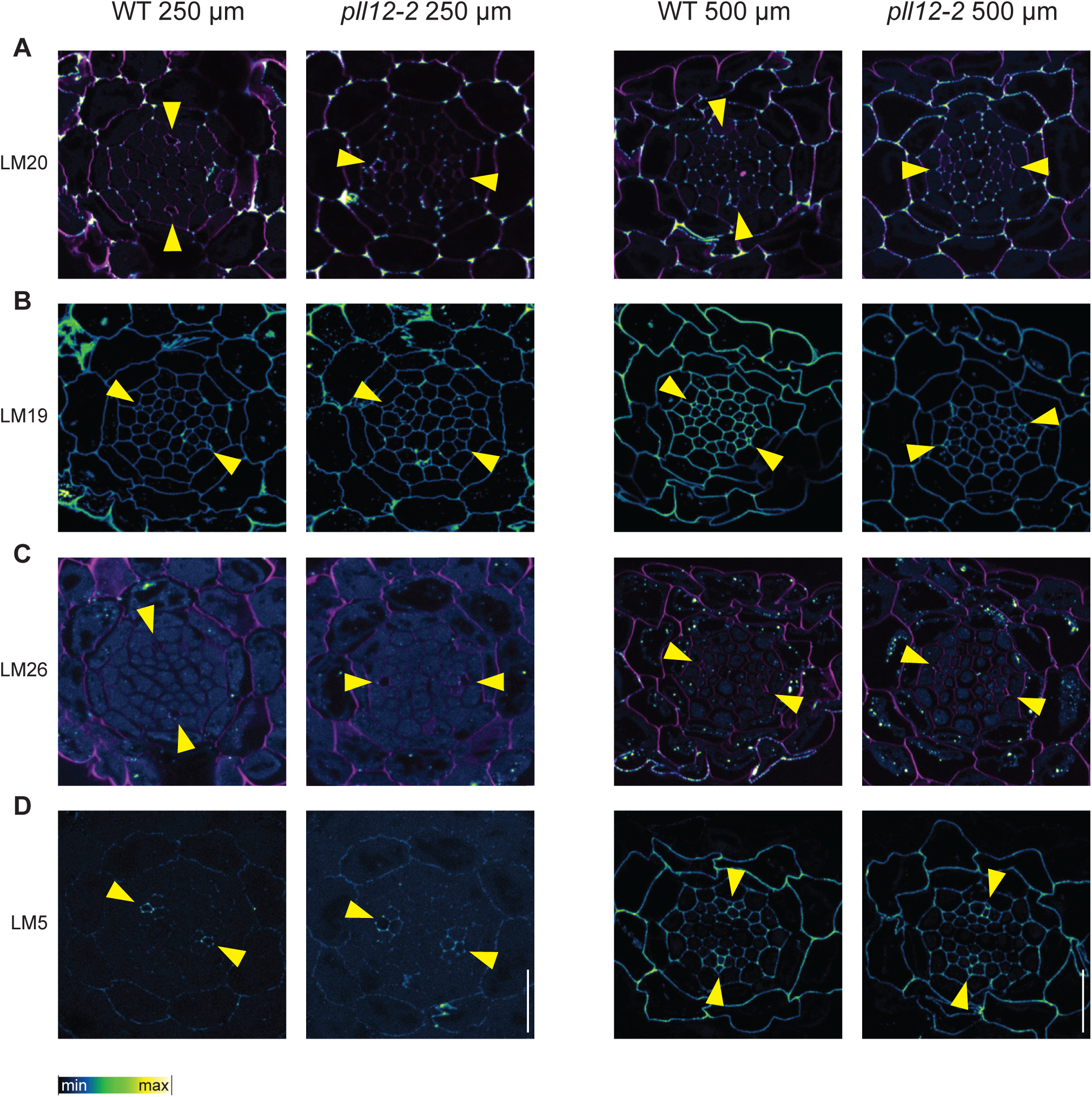
Pectin epitopes in root cross sections. **A-D**) Cross sections of 5-day-old roots at 250 μm (time of SE enucleation) and 500 μm (around time of MSE differentiation) A) LM20 detects a highly methyl-esterified pectin epitope. B) LM19 recognizes highly de-methylated pectin epitope. C)LM26 detects a branched galactan epitope. D) LM5 detects a linear galactan. Yellow arrowheads indicate position of phloem pole. Antibodies are presented in LUT Green Fire Blue; cell wall counterstaining depicted in magenta in (A) and (C) is Calcofluor White. Scale bar represents 20 μm (A-D).

**Supplementary Figure 5:**
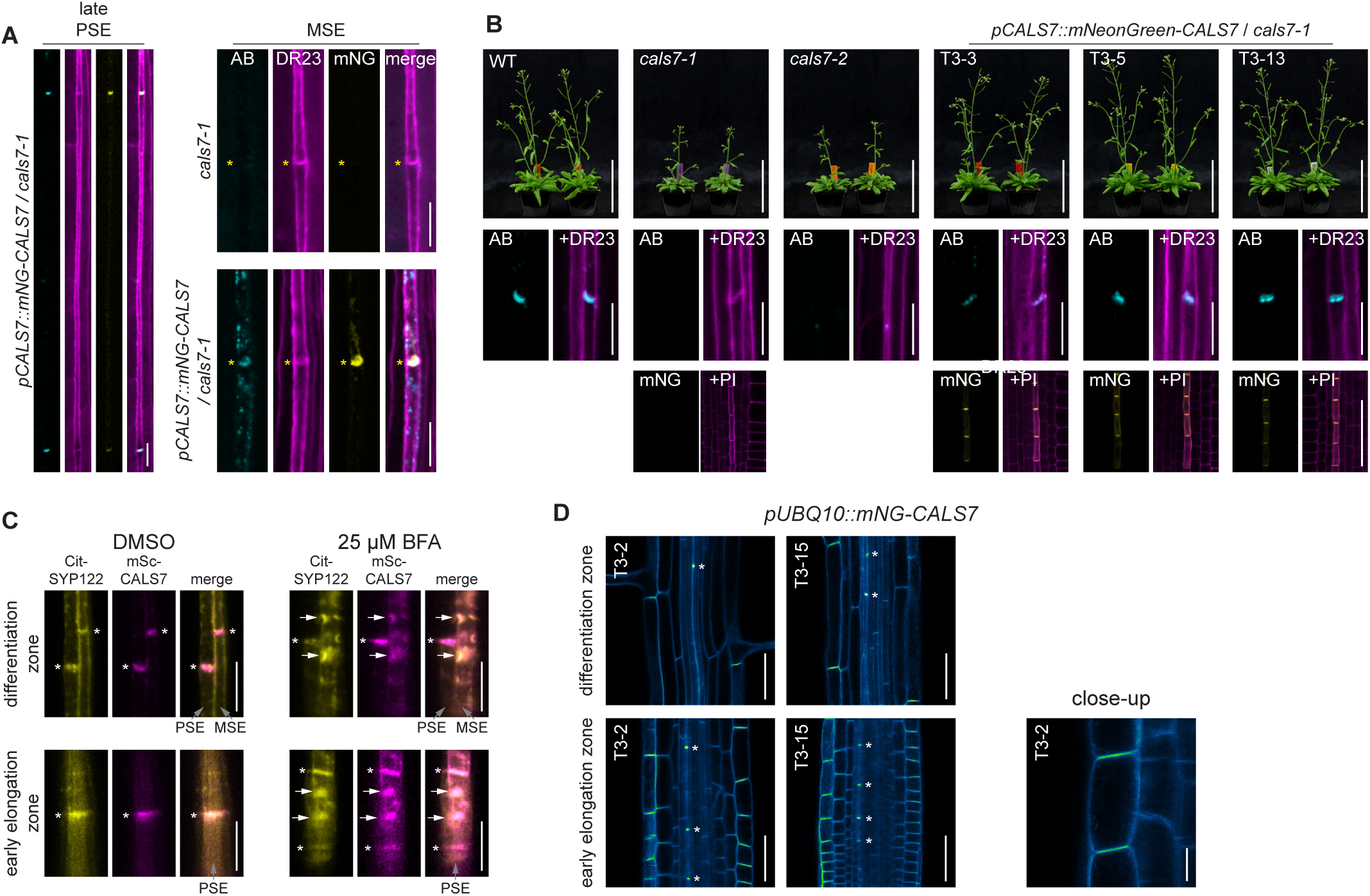
mNG-CALS7 is functional, sieve plate abundant and becomes immobilized at plasma membrane in a SE-specific manner. **A**) 5-day-old ClearSee-treated roots showing co-labeling of complementing mNG-CALS7 (yellow) under its endogenous promoter with Direct Red 23 (DR23, cell wall, magenta) and Aniline Blue (AB, callose, cyan) in fully differentiated PSE and MSE. **B**) Phenotypes of 2 mutant alleles and three independent complemented lines with respect to adult growth (42 dpg, top panel); in chemically fixed 5-day-old roots showing sieve plate callose (AB callose staining, cyan) counterstained with DirectRed 23 (magenta) in middle panel, and live-imaged mNG-CALS7 localization (yellow) in early PSE of 5-day-old roots counterstained with PI (magenta) in bottom panel. **C**) Sub-cellular localization of *pCASL7::mSc-CALS7* (magenta) and plasma membrane marker *pSUS6::Citrine-SYP122* (yellow) in young PSE (bottom panel) and late PSE next to young MSE (top panel) of 5-day-old roots in DMSO control (left) and upon 16h 25 μM BFA treatment (right). **D**) Live-imaging of ectopic *pUBQ10::mNG-CALS7* expression in early (bottom panel) and differentiating 5-day-old root (top panel) with close-up on early elongating epidermal cell (far-right), color coded for fluorescence intensity with LUT Green Fire Blue. Scale bars represent 10 μm in (A, B middle panel, C, D bottom right “close-up”), 100 mm (B top panel), and 50 μm (B bottom panel).

**Supplementary Figure 6:**
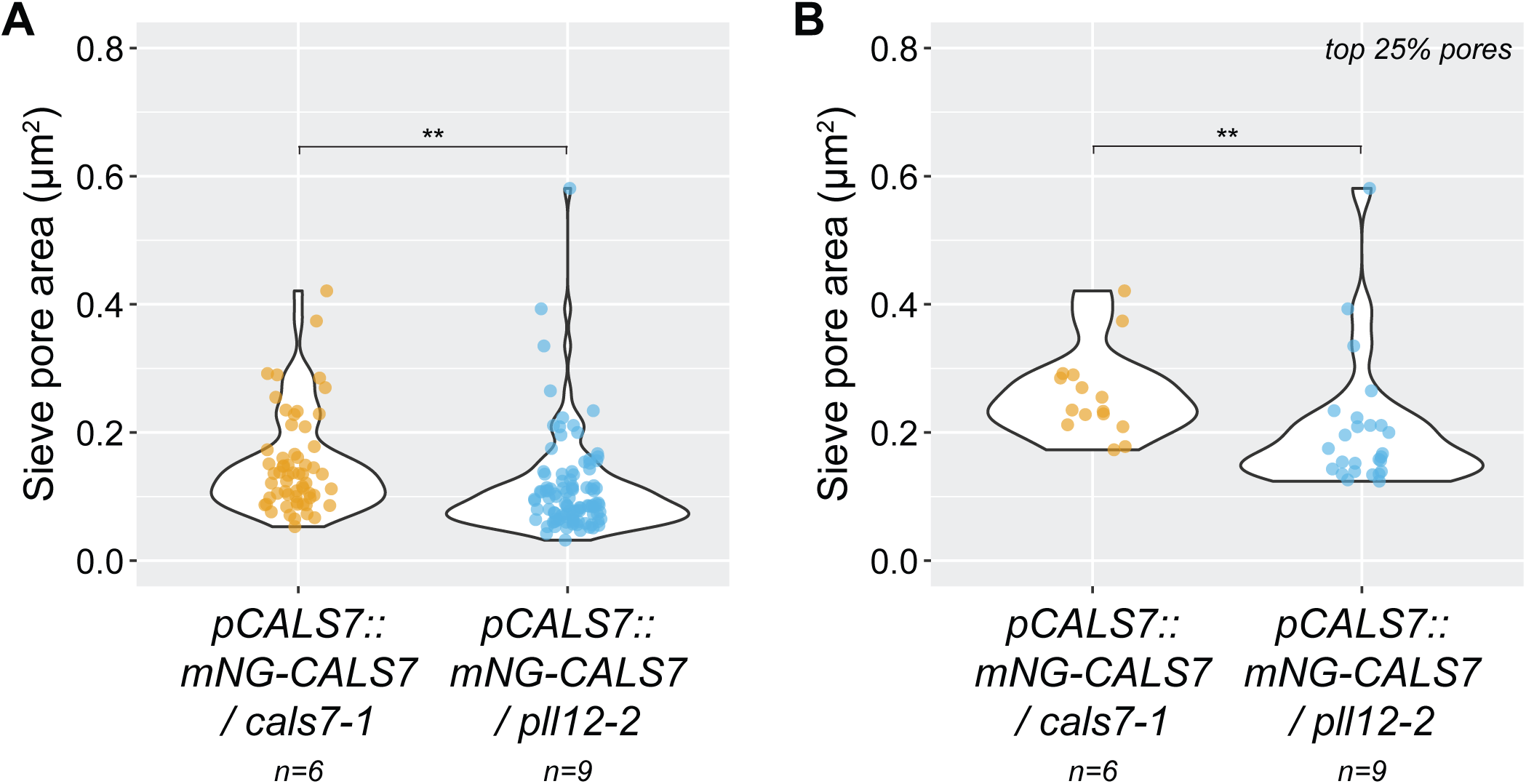
Sieve pore defects in *pll12* leaves. **A**-**B**) Quantification of sieve pore area in 21-day-old leaf vasculature presented in violin plots with overlaid dot plots considering all (A) and only the largest 25% of each genotype (B). Statistical significance was determined using a Wilcoxon sum rank test, significance levels: ** = *P* < 0.05. Raw data and p-values are presented in **Table S6**.

Supplementary Video 1: Time-lapse of root growth over 15 days for WT, *pll12-2, pPLL12::PLL12gen* / *pll12-2* and *pNEN4::PLL12cds* / *pll2-2* complemented lines; images assembled to video from scans for root growth assay (Fig. 1B, Fig. S1G) with one frame/day. Scale bar represents 50 mm.

Supplementary Table 1: Raw data, *P*-values and statistical grouping for ANOVA/Tukey tests on root growth (relates to Figure 1B and Supplementary Figure 1G).

Supplementary Table 2: Raw data, Statistical analysis, *P*-values and statistical grouping for Root dry biomass (relates to Fig. SI)

Supplementary Table 3: Raw data and statistical analysis for pSUC2::GFP fluorescence in root tip; relates to Fig. 3C.

Supplementary Table 4: Raw data, statistical analysis, P-values and statistical grouping for CFDA transport assay; relates to Fig. 3E.

Supplementary Table 5: Raw data, statistical analysis, and significance values for sieve pore areas in hypocotyls and leaf petioles; relates to Fig. 5C-D and Fig. S6A-B.

